# Centrosome-centromere capture range, rather than centrosome arrangement, determines multipolar chromosome segregation pattern after whole-genome duplication

**DOI:** 10.64898/2026.08.24.746909

**Authors:** Masaya Inoko, Guang Yang, Yuki Tsukada, Ryota Uehara

**Author notes:** Corresponding author: Ryota Uehara, Faculty of Advanced Life Science, Hokkaido University, Kita 21, Nishi 11, Kita-Ku, Sapporo 001-0021, Japan.

## Abstract

Whole-genome duplication (WGD) causes chromosome instability through multipolar chromosome segregation driven by supernumerary centrosomes. WGD cells formed through distinct processes, mitotic slippage (MS) and cytokinesis failure (CF), show a prominent difference in viability after multipolar chromosome segregation: MS causes a more skewed homologous chromosome distribution than CF, resulting in more frequent nullisomic chromosome segregation with poorer survival through the first mitosis. However, the determinants of route-dependent differences in post-WGD cell viability remain largely unknown, particularly regarding the contribution of spatial rearrangement of supernumerary centrosomes. Here, we found marked differences in supernumerary centrosome distribution upon entry into the first mitosis after MS and CF, stemming from distinct nuclear geometry. The distinct centrosome distributions differentiated kinetochore capture patterning after MS and CF, whereas their modulations had minimal effect on the fidelity of subsequent chromosome segregation. In contrast, artificially extending the centrosome-centromere capture range by depleting the microtubule depolymerizer MCAK drastically suppressed the MS-linked aggravation of nullisomic chromosome segregation through equalizing chromosome capture by each supernumerary centrosome. These results suggest that centrosome-centromere capture range, rather than the spatial arrangement of the centrosomes themselves, determines the fidelity of chromosome segregation after WGD. Our findings provide fundamental insights into atypical cell proliferation mechanisms after WGD.

**Summary statement:** Multipolar chromosome segregation patterns after whole-genome duplication are determined by centrosome-centromere capture range rather than supernumerary centrosome positioning.

## Introduction

Whole-genome duplication (WGD), which refers to the doubling of the entire set of chromosomes, contributes to broad biological phenomena including development, aging, evolution, and cancer (Conway et al., 2024; Darmasaputra et al., 2024; Tanaka et al., 2018; Van de Peer et al., 2017). WGD occurs in more than 30% of solid tumors (Bielski et al., 2018; Carter et al., 2012; Zack et al., 2013), fueling tumor heterogeneity by promoting chromosome instability (Dewhurst et al., 2014). A primary source of WGD-associated chromosome instability is multipolar chromosome segregation, induced by supernumerary centrosomes arising upon WGD. While most progenies formed through multipolar chromosome segregation are unviable, a small fraction survives and contributes to the promotion of aneuploidy and chromosome instability in the post-WGD cell populations (Duncan et al., 2012; Telentschak et al., 2015; Yang et al., 2012; Yang et al., 2026; Zhang et al., 2016). However, the determinants of survival and the fates of post-WGD progeny, particularly after multipolar chromosome segregation, remain largely unknown.

Mitotic slippage (MS-WGD, hereafter) and cytokinesis failure (CF-WGD) are two major causes of WGD in cancer cells (Was et al., 2022). MS-WGD is premature mitotic exit without proper chromosome segregation, occurring due to attenuation of the spindle assembly checkpoint (Mondal et al., 2007; Schnerch et al., 2013; Sze et al., 2004; Wang et al., 2002) or Cyclin B-Cdk1 activity (Brito and Rieder, 2006; Haschka et al., 2018; Sakurikar et al., 2012). CF-WGD often occurs through oncogene-mediated suppression of cytokinesis factors (Darp et al., 2022; Soeda et al., 2013) or chromosome segregation errors induced by DNA damage (Ganem and Pellman, 2012). Recent studies show that distinct WGD “routes” differentiate gene regulation and the fates of post-WGD progeny across organisms (Gemble et al., 2026; Rijnberk et al., 2022). Moreover, we recently found a prominent difference in multipolar chromosome segregation patterns and subsequent viability between MS- and CF-WGD cells, which was attributed to their distinct homologous chromosome arrangements (Inoko et al., 2026). After MS, homologous chromosomes show a skewed distribution within a nucleus, creating a large bias in the efficiency of centrosome-mediated homologous chromosome capture and promoting lethal nullisomic daughters in the subsequent mitosis (Inoko et al., 2026). In contrast, a more even distribution of homologous chromosomes after CF-WGD allows all centrosomes to capture them with less bias, resulting in much less nullisomic chromosome segregation. (Inoko et al., 2026).

Besides homologous chromosome distribution, spatial arrangement of centrosomes may influence multipolar chromosome segregation patterns after WGD. In normal diploid cells, two centrosomes, connected by linker proteins during interphase, separate and move apart during prophase (Remo et al., 2020). Eg5, a mitotic motor protein, primarily drives centrosome separation during prophase by mediating antiparallel microtubule sliding (Yildiz, 2025). Impaired prophase centrosome separation alters the geometric relationship between centrosomes and sister chromatids at mitotic entry, increasing the risk of chromosome segregation errors (Ganem et al., 2009; Kaseda et al., 2012; Silkworth and Cimini, 2012; Silkworth et al., 2009; Silkworth et al., 2012). In this study, we investigated the spatial rearrangement of supernumerary centrosomes during the first mitosis after MS- or CF-WGD, focusing on factors that affect lethal nullisomic chromosome segregation under different WGD conditions.

## Results and discussion

### Spatial distribution of the supernumerary centrosomes differs between MS- and CF-WGD cells in the first mitosis

We conducted live imaging to examine supernumerary centrosome dynamics during the first mitosis after pharmacological induction of MS- or CF-WGD in HCT116 cells (Inoko et al., 2026; see Materials and methods for details; Fig. 1A and B). Chromosomes and the centrosomes were labeled by stable expression of H2B-mCherry and EGFP-GCP3. Approximately 25% of cells underwent WGD in both MS- and CF-WGD conditions (Fig. S1A). We analyzed the remaining non-WGD diploids as an inner control. In the diploid control, 2 centrosomes, which were clustered with each other during interphase, became separated (over >4 μm at 26 ± 12 min before nuclear envelope breakdown (NEBD) (mean ± s.d., *n* = 18 cells from five independent experiments; Fig. S1B). Inter-centrosomal distance in diploid control kept increasing during prophase and reached 9.2 ± 3.8 μm at NEBD (mean ± s.d., *n* = 18 centrosome pairs from 18 cells from five independent experiments; Fig. 1C and D). Either in MS- or CF-WGD cells, the 4 centrosomes gained upon WGD were also clustered during the first interphase (Fig. 1B). These centrosomes became separated (beyond a circle with a 4 μm diameter) at 43 ± 23 or 39 ± 19 min before NEBD in MS- or CF-WGD cells, respectively (mean ± s.d., at least *n* = 20 cells from five independent experiments; Fig. S1B). After separation, centrosomes moved apart with heterogeneous kinetics, with some centrosomes moving farther while others moved only a short distance within single cells (Fig. S1C and D). However, the overall centrosome separation efficiency was equivalent between MS- and CF-WGD cells, with an average inter-centrosome distance at NEBD of 10.3 ± 3.1 and 10.2 ± 3.0 μm, respectively (mean ± s.d., at least *n* = 120 centrosome pairs from 20 cells from five independent experiments; Fig. 1C, D, and S1D).

**Figure 1:**
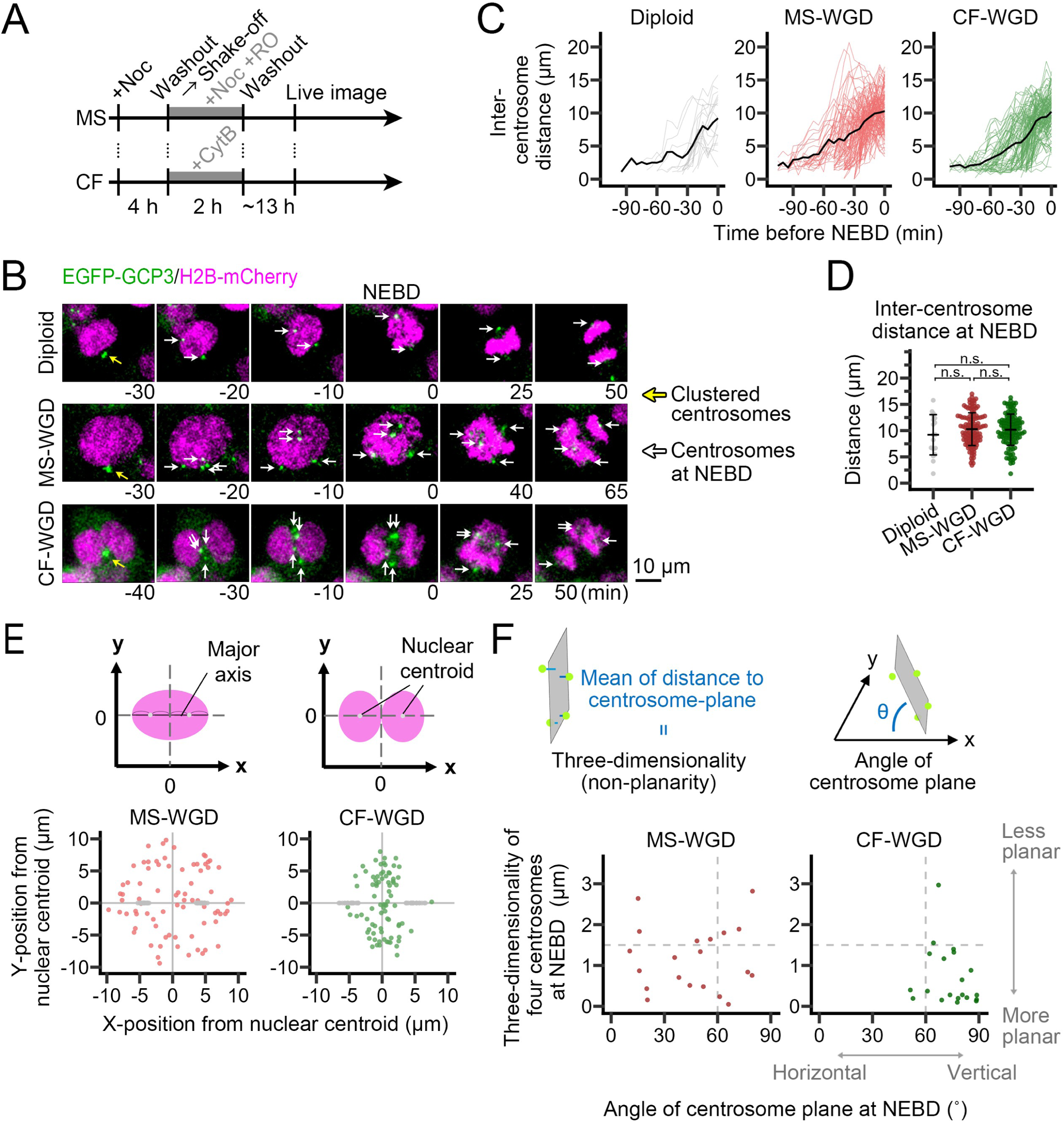
Distinct distribution of the supernumerary centrosomes upon the first mitotic entry between MS- and CF-WGD cells. **(A)** A scheme of WGD induction. **(B)** Time-lapse images of H2B-mCherry and EGFP-GCP3 throughout the first mitosis of diploid, MS-WGD, or CF-WGD cells. Images were taken at 5-min intervals. The NEBD timing is set to 0 min. Yellow arrows indicate clustered centrosomes before the onset of the first mitosis. White arrows indicate individual centrosomes after their separation. **(C)** Time-course of inter-centrosome distance during prophase (until NEBD) in B. Pooled data from at least 18 centrosome pairs from at least 18 cells from 5 independent experiments. **(D)** Inter-centrosome distance at NEBD in C. Mean ± s.e. from at least 5 independent experiments. There are no statistically significant differences between conditions (n.s.: not significant, the DSCF test). **(E)** Two-dimensionally projected positions of centrosomes relative to the nuclei at NEBD. As shown in the upper scheme, the x-axis was defined as the major axis of the fitted ellipse (for MS-WGD) or the line connecting the 2 nuclear centroids (for CF-WGD). At least 80 centrosomes from at least 20 cells from at least 5 independent experiments. Gray dots indicate nuclear positions in the scheme. **(F)** Three-dimensionality of 4 centrosomes (Y-axis) and angle of fitted centrosome plane (X-axis) in single WGD cells at NEBD in B. Schematic explanations of two indices are on the top (see Materials and methods for details). At least 20 cells from at least 5 independent experiments are analyzed.

We observed a striking difference in the spatial distribution of 4 centrosomes at NEBD between MS- and CF-WGD cells: Whereas 4 centrosomes showed a dispersed distribution around the nucleus in MS-WGD cells, all of them were trapped in the cellular equatorial region flanked by 2 nuclei in CF-WGD cells (Fig. 1B and Movie S1 and S2). Two-dimensional projections of centrosome positions relative to the centroids of the chromosome masses across multiple cells clearly recapitulate the difference in their spatial distributions between MS- and CF-WGD cells (Fig. 1E). To further evaluate the three-dimensional distribution of the centrosomes, we also measured the three-dimensionality (non-planarity) of 4 centrosomes and the orientation of the plane fitted to the 4 centrosomes (defined as “centrosome plane”) to the horizontal plane (Fig. 1F; note that a lower three-dimensionality indicates more planar arrangement of 4 centrosomes; see Materials and methods for detail). MS-WGD cells showed cell-to-cell heterogeneity in centrosomal three-dimensionality and orientation of the centrosome planes, indicating a polymorphic centrosome spatial arrangement in this condition (Fig. 1F and Movie S1). In contrast, centrosome arrangement was less heterogeneous (i.e., more consistent across cells), with lower three-dimensionality and vertical orientation in CF-WGD cells (Fig. 1F and Movie S2), indicating that the 4 centrosomes were distributed in a planar manner in the inter-nuclear region. As mitosis progressed, the lower centrosomal three-dimensionality in CF-WGD cells was gradually lost, particularly in the tetrapolar case, becoming less distinguishable from the MS-WGD condition by anaphase onset (AO) (Fig. S1E). Therefore, the difference in the centrosomal arrangement between MS- and CF-WGD cells became evident transiently for several minutes around NEBD.

### Spatial arrangement of kinetochore-microtubule attachment differs between MS- and CF-WGD cells in the first mitosis

The above difference in supernumerary centrosome distribution, albeit transient, may affect the subsequent kinetochore-microtubule attachment between MS- and CF-WGD cells. To address this possibility, we examined the distribution of kinetochores and spindle microtubules by immunostaining CENP-C and β-tubulin, respectively, around the NEBD of the first mitosis after MS- or CF-WGD (Fig. 2A; chromosomes were labeled with H2B-mCherry). In MS-WGD cells, most microtubules emanated from the spindle poles located outside the chromosome masses (Fig. 2A). These microtubules tended to attach to the kinetochores in a centripetal manner (i.e., from cell periphery to the center of the chromosome masses; Fig. 2A inset), with the captured kinetochores detected significantly farther from the centroid of chromosome mass than unattached kinetochores (Fig. 2B). In contrast, in CF-WGD cells, microtubules emanated from the spindle poles located close to the centroid of chromosome masses. These microtubules tended to capture the kinetochore in a centrifugal manner, with the captured kinetochores detected significantly closer to the centroid of the chromosome masses than unattached kinetochores (Fig. 2A, B and S2A). Therefore, reflecting transient differences in supernumerary centrosome distribution, the spatial arrangement of kinetochore capture varied between MS- and CF-WGD cells at early prometaphase.

**Figure 2.**
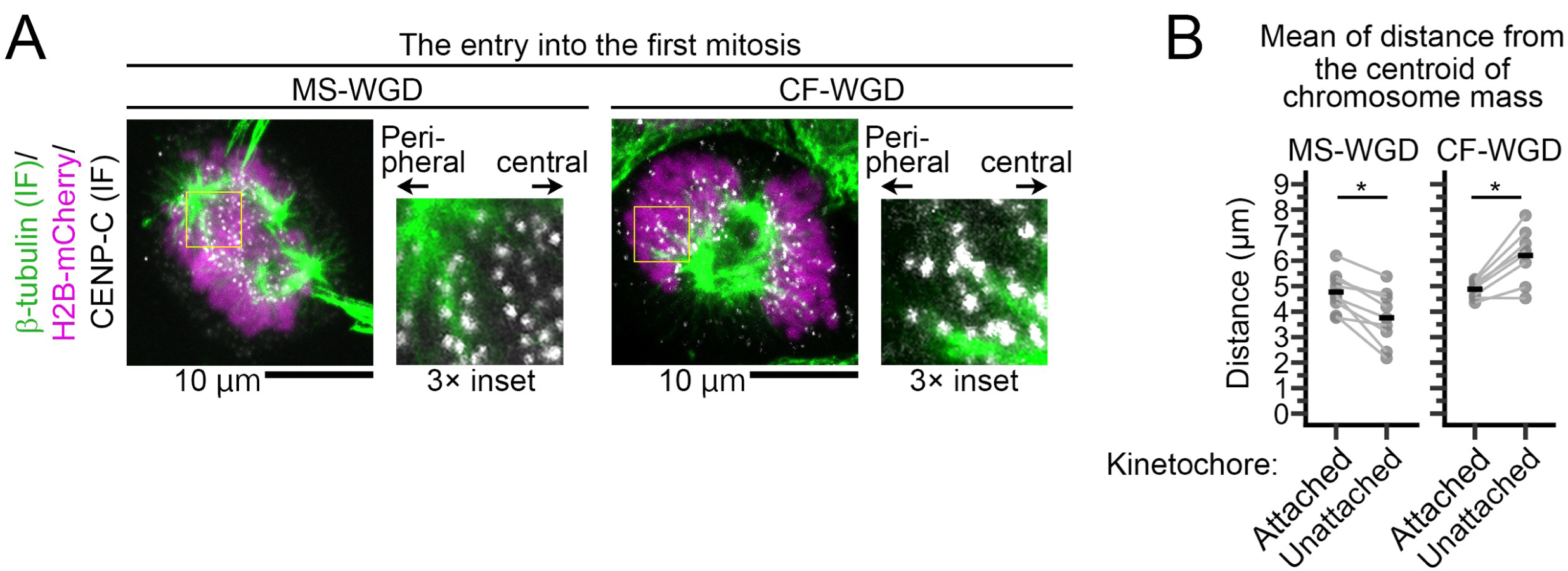
Distinct spatial patterns of kinetochore-microtubule attachment during the first mitosis between MS- and CF-WGD cells. **(A)** Fluorescence microscopy of H2B-mCherry, kinetochores (marked by CENP-C immunostaining), and microtubules (β-tubulin immunostaining) at the entry into the first mitosis after MS- or CF-WGD. Insets show 3x enlarged images. **(B)** Distance from the centroid of chromosome masses to kinetochores (attached or unattached by microtubules) in A. Individual data points represent mean values calculated from single cells (at least 8 cells from 3 independent experiments; \**p* < 0.05, the Welch’s t-test). Distributions of single kinetochore values are shown in Fig. S2.

### Forced centrosome separation in CF-WGD cells does not affect nullisomic chromosome segregation

We next addressed whether the noticeable difference in centrosome distributions contributed to the difference in chromosome segregation fidelity between MS- and CF-WGD cells (Inoko et al., 2026). For this, we sought to artificially modulate the characteristic centrosome arrangement around NEBD in CF-WGD cells. Since the characteristic planar centrosome arrangement was gradually lost after NEBD, it would have formed through physical hindrance or guidance of centrosome separation by the geometry of the 2 nuclei. Based on this idea, we tested whether enhancing centrosome separation through Eg5 overexpression (Eg5-OE) modulated centrosome arrangement in CF-WGD cells (Fig. 3A and B). Eg5-OE significantly increased the overall inter-centrosome distance at NEBD (Fig. 3C). Moreover, in CF-WGD cells, their centrosomes became distributed beyond the internuclear region in a pattern resembling that in MS-WGD cells (Fig. 3B and D). Consistently, the three-dimensionality of 4 centrosomes increased by Eg5-OE in CF-WGD cells (Fig. 3E). Eg5-OE modestly increased the inter-centrosome distance and three-dimensionality in MS-WGD cells (Fig. 3C and E).

**Figure 3.**
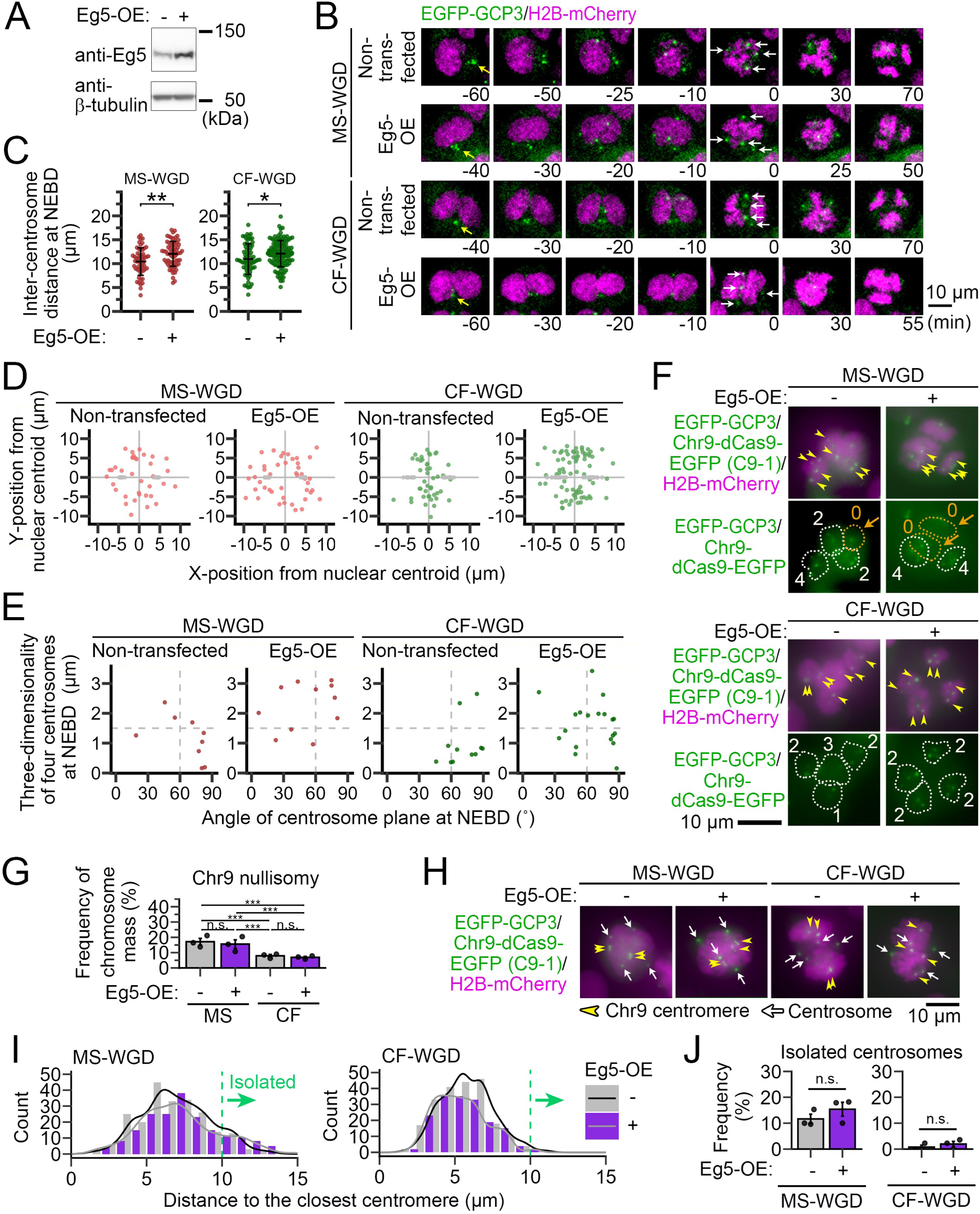
Effects of Eg5 overexpression on centrosome and centromere coordination after WGD. **(A)** Immunoblotting of Eg5 in non-transfected or Eg5-overexpressed (Eg5-OE) cells. β-tubulin was detected as a loading control. **(B)** Time-lapse images of H2B-mCherry and EGFP-GCP3 throughout the first mitosis of MS- or CF-WGD in non-transfected or Eg5-OE cells. Images were taken at 5-min intervals. The NEBD timing is set to 0 min. White arrows indicate centrosomes at NEBD. **(C)** Inter-centrosome distance at NEBD in B. Asterisks indicate statistically significant differences between conditions (\**p* < 0.05, \*\**p* < 0.01, the Welch’s t-test). **(D)** Two-dimensionally projected positions of centrosomes relative to the nuclei at NEBD. X- or Y-axis was defined as in Fig. 1G. At least 36 centrosomes from at least 9 cells from 5 independent experiments. Gray dots indicate the nuclear position in the scheme. **(E)** Three-dimensionality of 4 centrosomes (Y-axis) and angle of fitted centrosome plane (X-axis) in single WGD cells at NEBD in B. At least 9 cells from 5 independent experiments are analyzed. **(F)** Anaphase cells during tetrapolar division at the first mitosis after MS- or CF-WGD in non-transfected or Eg5-OE cells. The centrosomes, centromeres of chromosome 9, and chromosomes were labeled by EGFP-GCP3, dCas9-EGFP with specific sgRNA (C9-1), and H2B-mCherry, respectively. Yellow arrowheads indicate C9-1 foci. White or orange broken lines indicate chromosome masses with or without any C9-1 foci, respectively. Orange arrows indicate chromosome masses devoid of the labeled homologous chromosomes (nullisomy). The number of C9-1 foci in each chromosome mass is indicated in the bottom panels. **(G)** Frequency of C9-1 nullisomic chromosome segregation during tetrapolar anaphase at the first mitosis after MS- or CF-WGD induction in non-transfected or Eg5-OE cells. Means ± s.e. of three independent experiments. At least 240 segregated chromosome masses from at least 60 cells from 3 independent experiments were analyzed (n.s.: not significant, the Steel-Dwass test). **(H)** Cells at the entry into the first mitosis after MS- or CF-WGD in non-transfected or Eg5-OE cells. Each cell structure is labeled as in Fig. 3F. **(I)** Histogram of the distance from each centrosome to its closest C9-1 focus in H. Four centrosome-centromere distances were obtained from an individual cell. The centrosome-centromere combinations whose distances exceeded 10 μm (defined as “isolated centrosomes”) are indicated by the green arrow. At least 168 centrosome-centromere combinations from at least 42 cells from 3 independent experiments were analyzed. **(J)** Frequency of isolated centrosomes in I. Means ± s.e. of three independent experiments (n.s.: not significant, the Brunner-Munzel test).

We reasoned that if the characteristic difference in supernumerary centrosome arrangement contributed to the distinct prevalence of nullisomic chromosome segregation between MS- and CF-WGD cells (Inoko et al., 2026), then Eg5-OE-mediated modulation of centrosome arrangement would affect the process. Therefore, we analyzed the patterns of multipolar chromosome segregation of homologous chromosome 9 labeled by dCas9-EGFP co-expressed with a specific single-guide RNA (sgRNA, C9-1) (Ma et al., 2015) and EGFP-GCP3 labeling the centrosomes in anaphase or telophase during the first mitosis after MS- or CF-WGD in non-transfected or Eg5-OE cells (Fig. 3F) (Inoko et al., 2026). While around 30% of WGD cells in the non-transfected condition underwent tripolar chromosome segregation by clustering 2 of 4 centrosomes during prometaphase, this clustering was largely suppressed in Eg5-OE WGD cells, with the majority (more than 84%) undergoing tetrapolar chromosome segregation (Fig. S3A). For this reason, we analyzed the C9-1 segregation pattern during tetrapolar chromosome segregation (Fig. 3G and S3B). Consistent with the previous observation (Inoko et al., 2026), MS-WGD cells showed a significantly higher frequency of nullisomic C9-1 segregation compared to CF-WGD cells in the non-transfected condition (Fig. 3G and S3B). Despite drastic changes in centrosomal distribution by Eg5-OE, the frequency of nullisomic chromosome segregation was equivalent between non-transfected and Eg5-OE conditions either after CF-WGD or MS-WGD (Fig. 3G and S3B). These data indicate that the characteristic spatial arrangements of supernumerary centrosomes at mitotic entry, especially after CF-WGD, do not influence the prevalence of nullisomic chromosome segregation during subsequent multipolar division.

We previously found that the probability for a centrosome to fail to capture any homologous centromeres was highly correlated to the distance between the centrosome and its closest homologous centromere at NEBD (Inoko et al., 2026). Therefore, we tested whether Eg5-OE affected the centrosome-homologous centromere distance at NEBD in MS- or CF-WGD cells (Fig. 3H and I). Consistent with the previous observation, MS-WGD cells had longer centrosome-centromere distances than CF-WGD cells (on average 7.0 ± 2.3 or 5.8 ± 1.7 μm in non-transfected MS- or CF-WGD cells, respectively; mean ± s.d., at least *n* = 216 centrosome-centromere combinations from at least 54 cells from three independent experiments; Fig. 3I), with more frequent “isolated centrosomes,” located extremely far (>10 μm) from any of the C9-1 foci (Fig. 3J). Frequency of isolated centrosomes did not significantly change upon Eg5-OE either in MS- or CF-WGD cells (Fig. 3I and J), demonstrating the minimal impacts of supernumerary centrosome distribution on the spatial coordination between individual centrosome-homologous centromere pairs. These data collectively exclude the possibility that distinct arrangements of supernumerary centrosomes between MS- and CF-WGD cells contribute to their distinct prevalence of nullisomic chromosome segregation, aside from the influence of centrosome-centromere distance.

### MCAK depletion suppresses the nullisomic chromosome segregation after WGD

Our previous and current results support the idea that centrosome-centromere distance, rather than spatial arrangement of the centrosomes themselves, is the primary determinant of the prevalence of nullisomic chromosome segregation: When the distance to all homologous chromosomes from a certain centrosome exceeds the maximum distance range that microtubules from the centrosome can reach, that centrosome fails to capture and segregate any of these homologs. To test this idea, we attempted to extend the reach range of centromere capture by depleting the microtubule depolymerase MCAK (Hunter et al., 2003; Walczak et al., 2013) and examined its effects on nullisomic chromosome segregation in MS-WGD cells with highly skewed homolog distributions (Fig. 4A) (Inoko et al., 2026). MCAK depletion significantly reduced the frequency of nullisomic C9-1 segregation in MS-WGD cells in both tripolar and tetrapolar division (Fig. 4B and C). To further test whether MCAK depletion indeed reduced nullisomic C9-1 segregation through the extension of the reach range of centromere capture, we directly analyzed the relationship between centrosome-C9-1 homolog distance at NEBD and the frequency of subsequent capture of the C9-1 by the centrosome in live imaging (Fig. 4D and E) (Inoko et al., 2026). In the mock-depleted control, the frequency of C9-1 capture dropped sharply when the initial centrosome-C9-1 distance exceeded 10 μm (Fig. 4E), consistent with our previous observation (Inoko et al., 2026). MCAK depletion significantly promoted C9-1 capture over an initial centrosome-C9-1 distance of >10 μm (Fig. 4E), demonstrating that MCAK depletion indeed extended the reach range of centromere capture from each centrosome. We further analyzed the relationship between the distance from the centrosome to its closest C9-1 at NEBD (centrosome-closest C9-1 distance) and the frequency of centrosomes failing to capture any C9-1 foci (i.e., nullisomic chromosome segregation). Though centrosome-closest C9-1 distance slightly increased after MCAK depletion (Fig. 4F), the frequency of nullisomic chromosome segregation tended to decrease after MCAK depletion in the range of centrosome-closest C9-1 distance >10 μm (Fig. 4G and H). These results collectively suggest that the balance between centrosome-centromere distance and the range of centromere capture by the centrosome primarily determines chromosome segregation patterns during multipolar division, largely affecting the viability of post-WGD progeny undergoing early aberrant mitosis.

**Figure 4.**
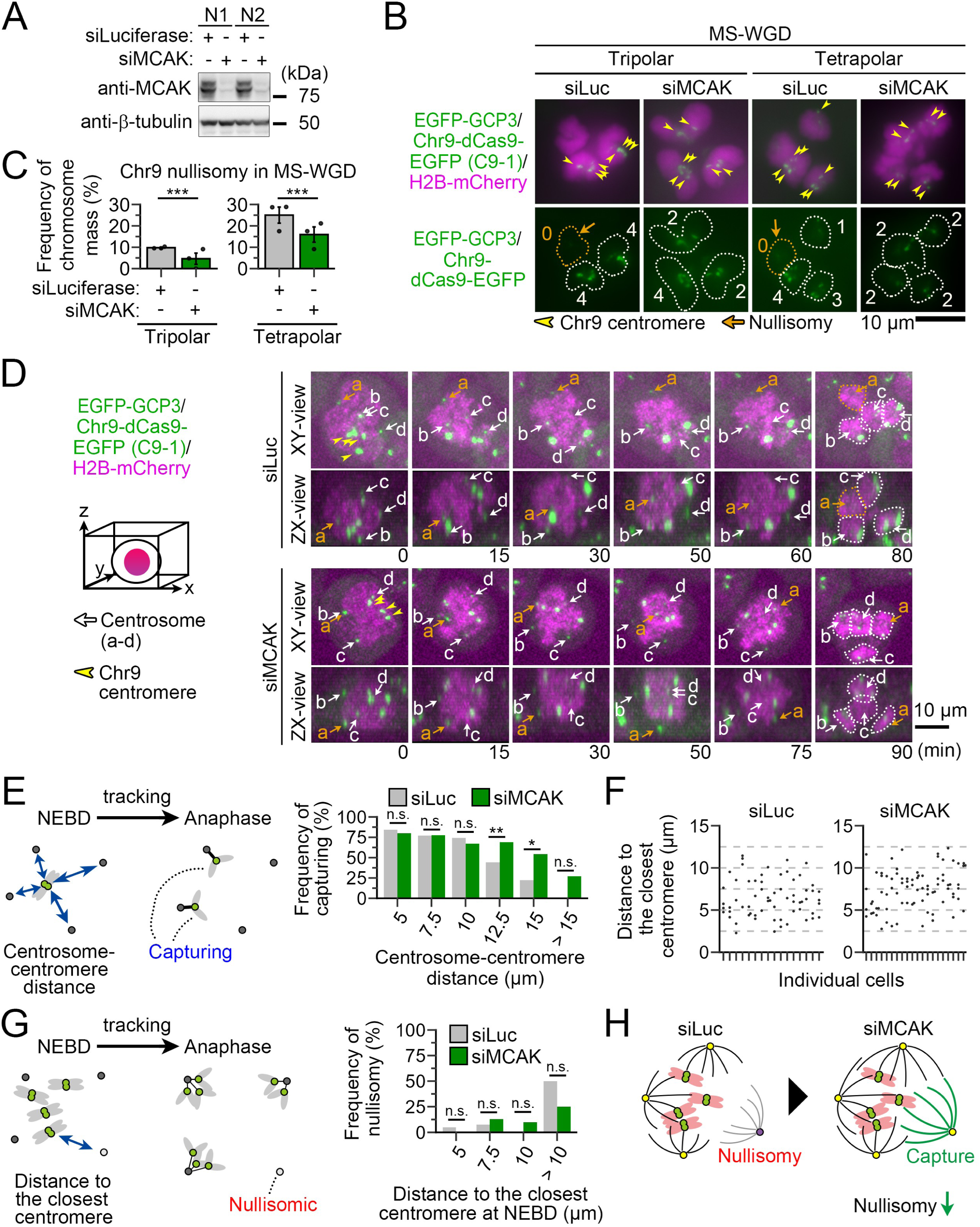
MCAK depletion suppresses subsequent nullisomic chromosome segregation. **(A)** Immunoblotting of MCAK in mock- or MCAK-depleted cells sampled at 18 h after WGD induction. β-tubulin was detected as a loading control. **(B)** Anaphase cells at the first mitosis after MS-WGD in mock- or MCAK-depleted conditions. Each cell structure is labeled as in Fig. 3F. Yellow arrowheads indicate C9-1 foci. White or orange broken lines indicate chromosome masses with or without any C9-1 foci, respectively. Orange arrows indicate chromosome masses devoid of the labeled homologous chromosomes (nullisomy). The number of C9-1 foci in each chromosome mass is indicated in the bottom panels. **(C)** Frequency of C9-1 nullisomic chromosome segregation during anaphase at the first mitosis after MS-WGD induction in mock- or MCAK-depleted cells. Means ± s.e. of 3 independent experiments. At least 81 segregated chromosome masses from at least 27 cells from 3 independent experiments were analyzed for each polarity. Asterisks indicate statistically significant differences between conditions (\*\*\**p* < 0.001, the Brunner-Munzel test). **(D)** Time-lapse images of H2B-mCherry, EGFP-GCP3, and dCas9-EGFP with C9-1 sgRNA throughout the first mitosis after MS-WGD in mock- or MCAK-depleted cells. Top and side views (XY and XZ planes, respectively) of the three-dimensional reconstructions are shown. Images were taken at 5-min intervals. The NEBD timing is set to 0 min. Individual centrosomes were distinguished by labeling different letters (a-d). White or orange arrows indicate centrosomes that are located ≤10 μm or >10 μm from the closest centromere at NEBD, respectively. White or orange broken lines indicate chromosome masses with or without any C9-1 foci, respectively. **(E)** The frequency of centrosome-centromere capturing, which was calculated for each bin of the initial centrosome-centromere distance in mock- or MCAK-depleted MS-WGD cells (schematized on the left). At least 256 centrosome-centromere pairs from 16 cells from 8 independent experiments were analyzed. **(F)** Quantification of the distance from each centrosome to its closest centromere at the entry into the first mitosis after MS-WGD in mock- or MCAK-depleted cells. Each column represents the distances obtained in an individual cell. At least 16 cells from 8 independent experiments were analyzed. **(G)** The frequency of the centrosomes that failed to capture any C9-1, which was calculated for each bin of the closest centrosome-centromere distance at NEBD (schematized on the left). At least 64 centrosomes from at least 16 cells from 8 independent experiments were analyzed. **(H)** A schematic model for determinants of (un) equality in chromosome capture upon multipolar chromosome segregation after WGD.

In this study, we investigated the dynamics of supernumerary centrosomes and chromosome segregation during the first mitosis after MS- or CF-WGD in HCT116 cells. Our results suggest that two aspects of centrosomal control, described below, contribute separately to the spatial organization of chromosome capture or segregation. Firstly, distinct supernumerary centrosome distribution upon mitotic entry, determined through nuclear geometry and level of Eg5 function, differentiates the early kinetochore capture orientation between MS- and CF-WGD cells (Fig. 2). However, this early difference in centrosome-kinetochore coordination would be gradually canceled, as centrosomes are subsequently redistributed toward poles during prometaphase (Fig. S1E). As a result, the initial arrangement of the supernumerary centrosomes has minimal influence on the later chromosome segregation pattern. Second, the balance between centrosome-centromere distance and centromere capture range determines the equality of chromosome segregation between poles during multipolar segregation, primarily affecting the prevalence of nullisomic chromosome segregation that characterizes early viability after WGD via different routes (Inoko et al., 2026). Our findings provide new insight into the principles underlying route-dependent diversification of cell fates following WGD.

## Materials and methods

### Cell culture and cell line establishment

HCT116 cells (WT line, RRID: CVCL_0291 from RIKEN BRC) were cultured in McCoy’s 5A (Wako) supplemented with 10% fetal bovine serum (FBS) and 1× antibiotic-antimycotic solution (AA; Sigma-Aldrich) at 37°C with 5% CO_2_. Transgenic cell lines were established by transfecting HCT116 cells with plasmids encoding the corresponding transgenes using JETPEI (Polyplus-transfection), followed by selection of positive clones with the appropriate antibiotics. Cell lines used in this study are listed in Table S1. All experiments were performed with mycoplasma-free cells (tested by MycoStrip, mycoplasma detection kit; InvivoGen).

### Plasmids, compounds, and antibodies

The plasmid vectors constructed or used in this study are listed with the primer information in Table S2. Compounds and antibodies used in this study are listed in Table S3. The siRNAs used in this study are 5’-GCAAUAAACCCAGAACUCUtt-3’ (MCAK; DNA in lowercase) and 5’-CGUACGCGGAAUACUUCGAtt-3’ (luciferase). siRNA transfection was performed using Lipofectamine RNAiMAX (Thermo Fisher Scientific).

### WGD induction

To induce WGD, cells were first arrested at prometaphase by treating them with 40 ng/mL nocodazole for 4 h, washed 3 times with supplemented culture medium, and then shaken off the culture dishes. For CF-WGD induction, we treated the shaken-off mitotic cells with 5 μg/mL cytochalasin B for 2 h. For MS-WGD induction, we co-treated the shaken-off mitotic cells with 40 ng/mL nocodazole and 10 μM RO3306 for 2 h. Throughout the manuscript, we defined the WGD induction time point as when we added these inhibitors to cell cultures. After 2 h of WGD induction, cells were washed 3 times with supplemented culture medium. The efficiency of WGD induction was evaluated as previously described, either by the presence of 2 nuclei (for CF-WGD) or by the increase in nuclear areas to >160 μm^2^ (for MS-WGD) (Inoko et al., 2026).

#### Live imaging

For live imaging of the first mitosis after WGD induction, we seeded WGD-induced cells on an 8-well cover glass-bottom chamber (zell-kontakt GmbH or IWAKI 5232-008), replaced culture media with supplemented phenol red-free McCoy’s 5A (Cytiva) at 14 h after WGD induction, and started cell imaging from 15 h after WGD induction.

#### Quantitative analysis of the centrosome positions in the first mitosis

To analyze the centrosome positions in Fig. 1 and 3, EGFP-GCP3/H2B-mCherry cells were imaged by live cell microscopy. We obtained centrosome coordinates using a semi-automated algorithm in FIJI. The ImageJ macro files used for centrosome foci detection are provided as Supplemental material S1. We automatically detected candidate centrosome coordinates and then manually curated them to identify the correct ones. The centroid of the mitotic chromosome masses was measured using FIJI from the auto-segmented regions of H2B-mCherry signals on the middle slice of Z-sections covering the entire chromosome masses.

To quantify the spatial arrangement of supernumerary centrosomes, a single plane was fitted to the 4 centrosomes in each WGD cell using the least-squares method (to obtain a centrosome plane). The MATLAB file used for plane fitting is provided as Supplemental material S2. Three-dimensionality (non-planarity) is defined as the mean of the perpendicular distances from the centrosome plane to each of the 4 centrosomes. Lower or higher values of three-dimensionality indicate more or less planar arrangements of the 4 centrosomes, respectively. The angle between the centrosome plane and the horizontal plane is calculated from the normal vector of the fitted plane. Angle values closer to 90° or 0° indicate a more vertically or horizontally oriented plane, respectively.

#### Quantitative analysis of the centrosome and C9-1 centromere positions in the first mitosis

To analyze the centrosome and C9-1 centromere positions in Fig. 4, EGFP-GCP3/dCas9-EGFP (C9-1)/H2B-mCherry cells were imaged by live cell microscopy. The positions of the centrosomes and C9-1 foci were manually measured using the cell counter tool in FIJI.

#### Immunostaining

Cells were fixed with 100% methanol at −20°C for 10 min, treated with BSA blocking buffer (150 mM NaCl, 10 mM Tris-HCl, pH 7.5, 5% BSA, and 0.1% Tween 20) for 30 min at 25°C, and incubated with the first antibodies overnight at 4°C and then with fluorescence-conjugated secondaries overnight at 4°C. Following each treatment, cells were washed 3 times with phosphate-buffered saline (PBS).

For the immunostaining analysis of kinetochore capture in early prometaphase, we analyzed only centromeres clearly identifiable as CENP-C foci. Among the observed CENP-C foci, those associated with microtubule signals were classified as attached, whereas those lacking detectable microtubule signals were classified as unattached. The centroid of the mitotic chromosome masses was measured using FIJI from the auto-segmented regions of H2B-mCherry signals on the middle slice of Z-sections covering the entire chromosome masses.

#### Microscopy

Fixed cells were observed under either of the following microscopes: A Ti2 microscope (Nikon) with a ×100 1.45 NA Plan-Apochromatic and Zyla4.2 sCMOS camera (Andor); a Ti2-E microscope (Nikon) with ×100 1.4 NA Plan-Apochromatic-VC and an A1 confocal microscope system (Nikon).

Live cell imaging was conducted under the following microscope at 37°C with 5% CO_2_: A TE2000 microscope with x60 1.4 NA Plan-Apochromatic, CSU-X1 (Yokogawa), and an iXon3 EMCCD camera (Andor). Image acquisition was controlled by μManager.

#### Immunoblotting

Cells were lysed in SDS/PAGE sample buffer (1.125% SDS, 35 mM Tris-HCl, pH 6.8, 11.25% glycerol, 5% 2-mercaptoethanol), boiled for 5 min, and subjected to SDS/PAGE. Separated proteins were transferred onto Immun-Blot PVDF membrane (Bio-Rad). The blotted membranes were blocked with 0.3% skim milk in Tween Tris-buffered saline (TTBS; 50 mM Tris, 138 mM NaCl, 2.7 mM KCl, and 0.1% Tween 20), incubated with the first antibodies for 1 or 2 h at 25 °C, or overnight at 4 °C, and incubated with horseradish peroxidase (HRP)-conjugated secondary antibodies for 1 or 2 h at 25 °C. Each step was followed by 3 washes with TTBS. We detected signals using ezWestLumi plus ECL Substrate (ATTO, Tokyo, Japan) with a LuminoGraph II chemiluminescent imaging system (ATTO).

#### Statistical analysis

Analyses for significant differences between the two groups were conducted using the two-tailed Welch’s *t*-test, the Fisher exact test, or the Brunner-Munzel test in R software (The R Foundation). Multiple group analyses were conducted using the Steel-Dwass test or the Dwass-Steel Critchlow-Fligner (DSCF) test in R software. Statistical significance was set at *p* < 0.05. *P*-values are indicated in figures or the corresponding figure legends.

## Acknowledgment

We are grateful to Renata Basto for valuable discussions, Kinya Yoda for antibodies, the NIC at Hokkaido University for microscopes, and the Information Initiative Center at Hokkaido University for the Large-scale Computing System. MI and GY are supported by JST SPRING, Grant Number JPMJSP2119. This work was supported by JSPS KAKENHI (Grant Numbers JP19KK0181, JP19H05413, JP19H03219, JPJSBP120193801, JP21K19244, JP22H04926, JP24K21956, JP24KK0139, and JP24K02017 to R.U.), the Princess Takamatsu Cancer Research Fund, the Kato Memorial Bioscience Foundation, the Orange Foundation, the Smoking Research Foundation, Daiichi Sankyo Foundation of Life Science, the Akiyama Life Science Foundation, the Hoansha Foundation, Sumitomo Electric Group CSR Foundation, and the Terumo Life Science Foundation to R.U. The authors declare no competing financial interests.

## Author Contributions

Conceptualization, M.I., and R.U.; Methodology, M.I., G.Y., Y.T., and R.U.; Investigation, M.I.; Formal Analysis, M.I.; Resources, M.I., G.Y., Y.T., and R.U.; Supervision, R.U.; Writing – Original Draft, M.I., and R.U.; Writing – Review & Editing, M.I., and R.U.; Funding Acquisition, M.I., and R.U.

## Data and resource availability

All relevant data and details of resources can be found within the article and its supplementary information.

**Table S1. List of cell lines used in this study**

**Table S2. List of plasmids used in this study**

**Table S3. List of antibodies and compounds used in this study**

**Supplemental movie S1: Three-dimensional views of centrosomes and chromosome mass at NEBD after MS-WGD**

Three-dimensionally constructed image of EGFP-GCP3 and H2B-mCherry (left) or EGFP-GCP3 (right) at NEBD after MS-WGD.

**Supplemental movie S2: Three-dimensional views of centrosomes and chromosome masses at NEBD after CF-WGD**

Three-dimensionally constructed image of EGFP-GCP3 and H2B-mCherry (left) or EGFP-GCP3 (right) at NEBD after CF-WGD.

**Supplemental material S1. The ImageJ macro and Python files used for the centrosome coordinates detection**

**Supplemental material S2. The MATLAB file used for the centrosome-plane fitting**

---

**Figure S1.**
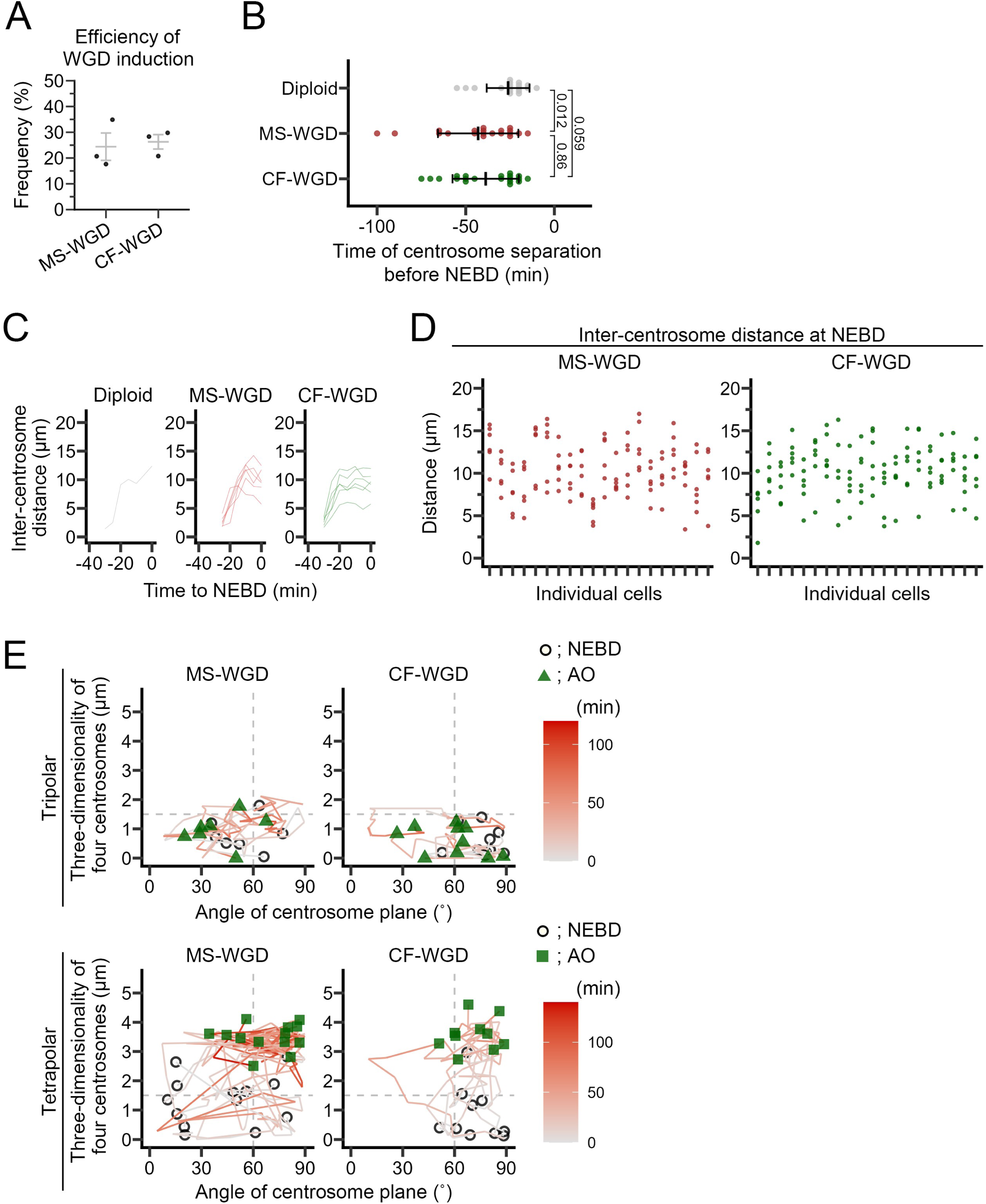
Live imaging analyses of the supernumerary centrosome dynamics during the first mitosis after MS- or CF-WGD. **(A)** Efficiency of WGD induction in Fig. 1B. Means ± s.e. of three independent experiments. At least 106 cells were analyzed in each condition. **(B)** The timing of centrosome separation before NEBD at the first mitosis after MS- or CF-WGD, or diploid inner-control in Fig. 1B. Means ± s.d. of at least 18 cells from at least 5 independent experiments. *p*-values obtained by the DSCF test between conditions are shown. **(C)** Time-course of inter-centrosome distance during prophase from representative single cells in Fig. 1B and C. **(D)** Quantification of inter-centrosome distance at NEBD in the first mitosis after MS- or CF-WGD in single cells. Each column represents the values acquired for every single cell. At least 20 cells from at least 5 independent experiments were analyzed. **(E)** Time-course of three-dimensionality of 4 centrosomes (Y-axis) and angle of fitted centrosome plane (X-axis) in single WGD cells. White circles and green triangles or squares (for tripolar or tetrapolar samples, respectively) indicate the value at NEBD or AO, respectively. The time-course changes in X and Y values are shown as color-coded line plots. At least 20 cells from at least 5 independent experiments are analyzed.

**Figure S2.**
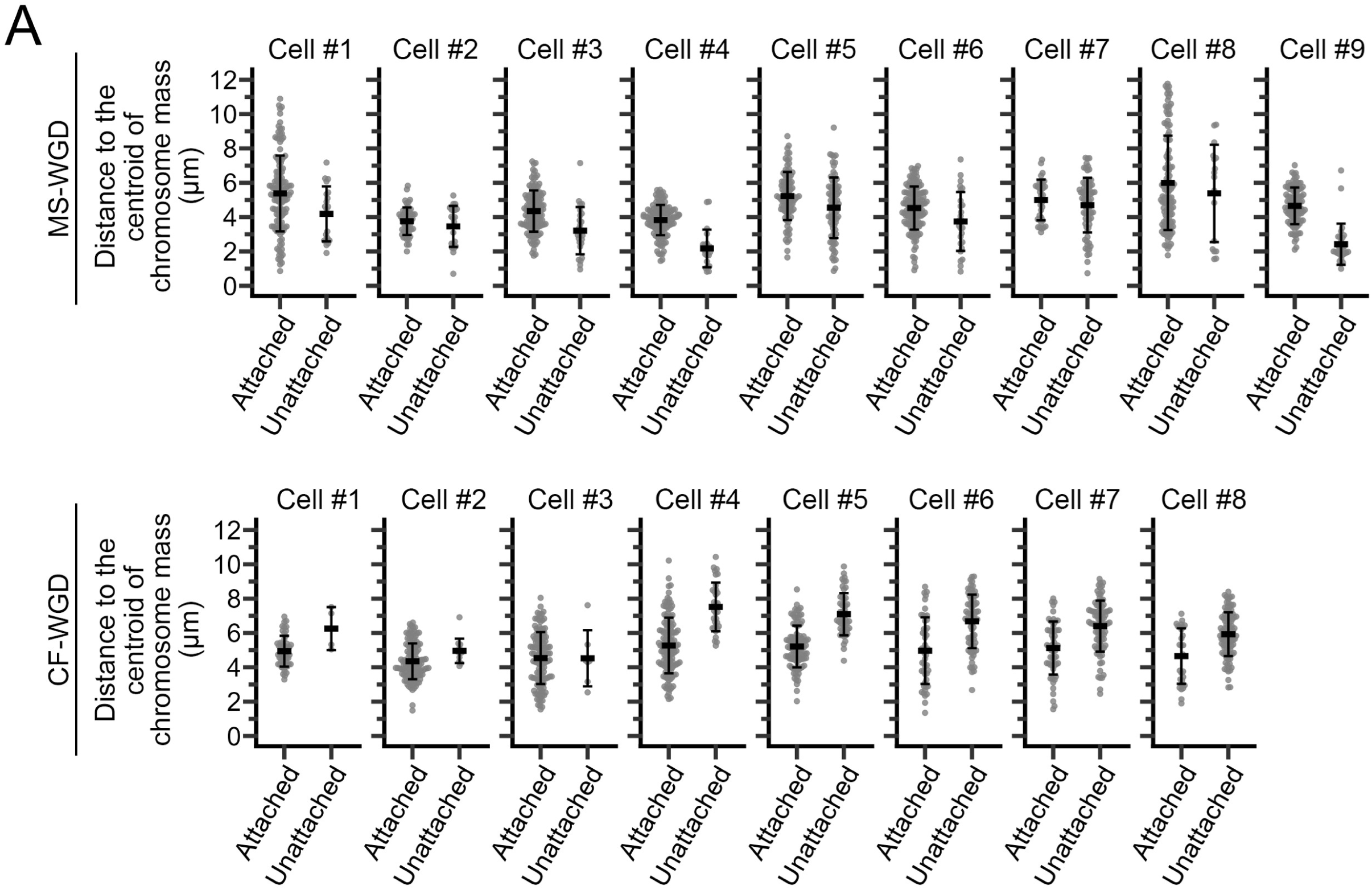
Imaging analysis of kinetochore-microtubule attachment patterns at the beginning of the first mitosis after WGD. **(A)** Distance from the centroid of chromosome masses to microtubule-attached and unattached kinetochores in individual cells in Fig. 2A. Means ± s.d. of at least 56 kinetochores are shown for every single cell.

**Figure S3.**
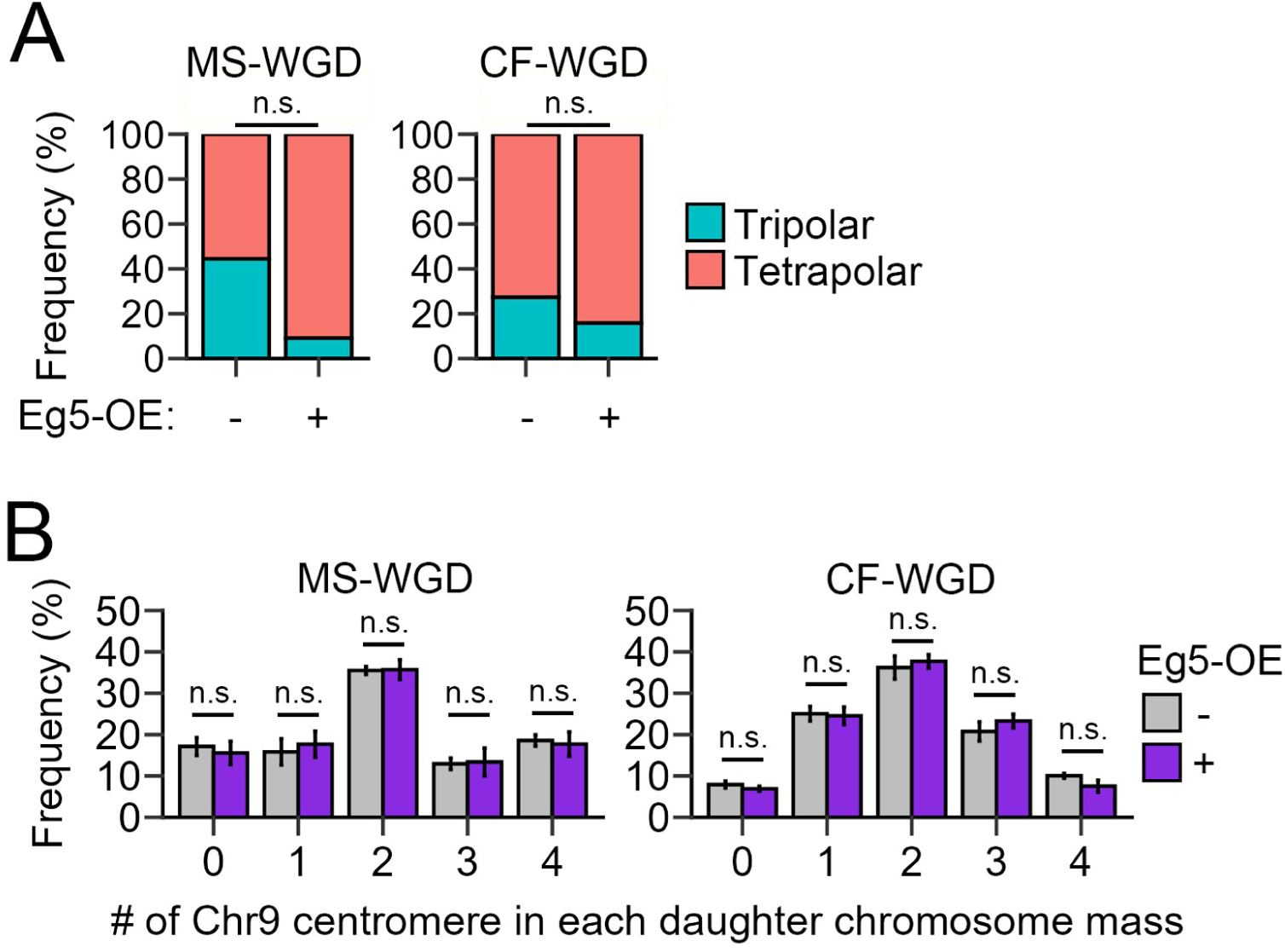
Effects of Eg5 overexpression on mitotic profiles after MS- or CF-WGD. **(A)** Frequency of chromosome segregation polarity at first mitosis after MS- or CF-WGD induction in non-transfected or Eg5-OE cells. At least 9 cells from 5 independent experiments were analyzed. The *p*-value from the Fisher exact test is 0.13 or 0.64 in MS- or CF-WGD, respectively. **(B)** Histograms of C9-1 foci numbers detected in segregated chromosome masses in anaphase in Fig. 3F. Means ± s.e. of 3 independent experiments. There are no statistically significant differences between MS- and CF-WGD in each category (n.s.: not significant, the Brunner-Munzel test).

**Figure S4.**
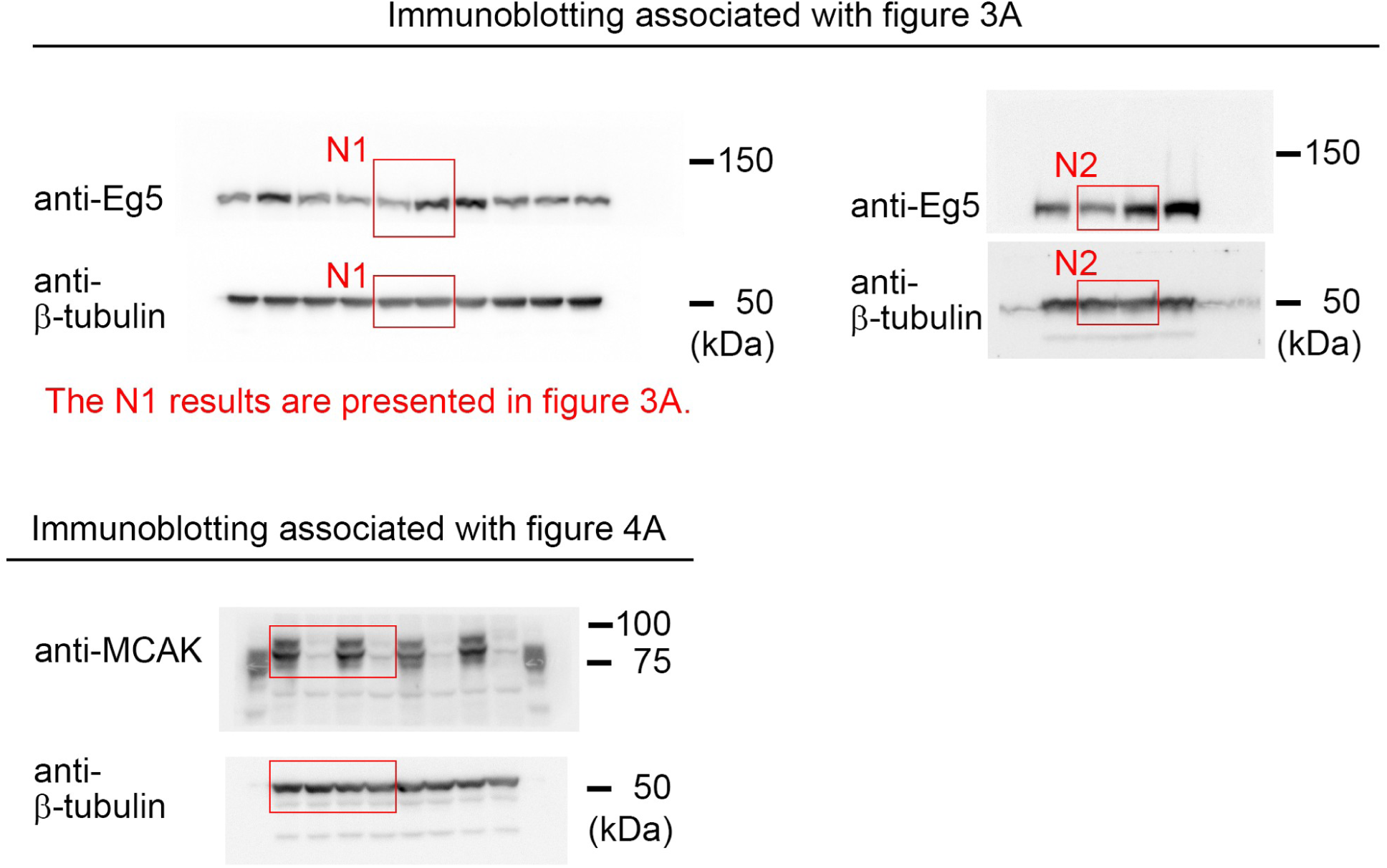
Original immunoblot membranes used for the analysis. Uncropped images of all immunoblots are shown, with the approximate indications of the cropped regions.

