## Supplementary MovieS1-S2, Supplementary Material S1-S2 for "Centrosome-centromere capture range, rather than centrosome arrangement, determines multipolar chromosome segregation pattern after whole-genome duplication": 220607_auto_select_check1.py

```
import pandas as pd 
import numpy as np
import glob
import re
import os
import argparse

#MaxResult = []
#MaxResult = pd.DataFrame()

list = glob.glob("F:/220708_12_21_29_0810_12_MI10_4N_prometa-anaphase/centrosome_analysis_data/result_of_auto_select_check1/221207/*")
for file in list:
    file1 = pd.read_csv(file)
    file2 = pd.DataFrame(file1)
    name = os.path.splitext(os.path.basename(file))[0]
    #print(name)

    ##列checkに1が入っている行を選択
    check = file2[file2["check"]==1]
    
    ##列checkに1が入っている行を別ファイルとして保存
    checkFile = pd.DataFrame(check)

    ##ファイルとして出力
    checkFile.to_csv("F:/220708_12_21_29_0810_12_MI10_4N_prometa-anaphase/centrosome_analysis_data/check_coordinates後check済ファイル/221207/{}_result.csv".format(name),index=False,encoding = "utf_8_sig")

##最大値を別ファイルに保存
#MaxFile = pd.DataFrame(MaxResult)
#MaxFile.to_csv("C:/Users/in0k0/Desktop/python/python勉強会/yamamotosan_result/max.csv", index=False)
```
