## Supplementary MovieS1-S2, Supplementary Material S1-S2 for "Centrosome-centromere capture range, rather than centrosome arrangement, determines multipolar chromosome segregation pattern after whole-genome duplication": check_coordinates.ijm

```
diam = 5;
Fold = 3;
FFold = 2; // fold for the z-axis of left and top projections
thresh_dist = 2;  // parameter for overlapping distance

showMessage("Select Open Folder");
openDir = getDirectory("Choose a Directory");
showMessage("Select Save Folder");
saveDir = getDirectory("Choose a Directory");

list = getFileList(openDir);
run("Set Measurements...", "mean centroid center stack redirect=None decimal=3");

for (i=0; i<list.length;i++){
    operation();
    display();


};


function display(){
	open(openDir+list[i]);
	n = getTitle();
	Stack.getDimensions(width, height, channels, slices, frames);
	wid = width*Fold;
	hei = height*Fold;
	wait(500);
	run("Size...", "width=wid height=hei time=frames constrain average interpolation=Bilinear");
	
	//    for Left projection
	wait(500);
	run("Reslice [/]...", "output=2.000 start=Left rotate avoid");
	left_w = getTitle();	
	run("Z Project...", "projection=[Max Intensity] all");
	Stack.setPosition(1,1,1);
	run("Enhance Contrast", "saturated=0.35");
	Stack.setPosition(2,1,1);
	run("Enhance Contrast", "saturated=0.35");
	left_p = getTitle();
	Stack.getDimensions(width, height, channels, slices, frames);
	widF = width*Fold*FFold;
	wait(500);
	run("Size...", "width=widF height=hei time=frames average interpolation=Bilinear");

	//    for top projection
	selectWindow(n);
	wait(500);
	run("Reslice [/]...", "output=2.000 start=Top avoid");
	top_w = getTitle();
	run("Z Project...", "projection=[Max Intensity] all");
	Stack.setPosition(1,1,1);
	run("Enhance Contrast", "saturated=0.35");
	Stack.setPosition(2,1,1);
	run("Enhance Contrast", "saturated=0.35");
	top_p = getTitle();
	Stack.getDimensions(width, height, channels, slices, frames);
	heiF = height*Fold*FFold;
	wait(500);
	run("Size...", "width=width height=heiF time=frames average interpolation=Bilinear");

	//    for hight projection
	selectWindow(n);
	run("Z Project...", "projection=[Max Intensity] all");
	Stack.setPosition(1,1,1);
	run("Enhance Contrast", "saturated=0.35");
	Stack.setPosition(2,1,1);
	run("Enhance Contrast", "saturated=0.35");
	n_p = getTitle();

	selectWindow(n);
	close();
	selectWindow(left_w);
	close();
	selectWindow(top_w);
	close();

	selectWindow(n_p);
	Stack.getDimensions(width, height, channels, slices, frames);
	
	setColor(255,255,255);
	NN = nResults;
	for(j=1;j<=NN-1;j++){
		if(getResult("Slice",j) != 0){
			selectWindow(n_p);
			Frame_n = getResult("Frame",j);
			Stack.setPosition(1,1,Frame_n);
			X_coordi = getResult("XM",j);
			Y_coordi = getResult("YM",j);
			X_coordiF = X_coordi*Fold;
			Y_coordiF = Y_coordi*Fold;
			drawOval(X_coordiF, Y_coordiF, diam, diam);
			drawString(j, X_coordiF+10, Y_coordiF);
			
			selectWindow(left_p);
			Stack.setPosition(1,1,Frame_n);
			S_coordi = getResult("Slice",j);
			S_coordiF = S_coordi*Fold*FFold;
			drawOval(S_coordiF, Y_coordiF, diam, diam);
			drawString(j, S_coordiF+10, Y_coordiF);
			
			selectWindow(top_p);
			Stack.setPosition(1,1,Frame_n);
			drawOval(X_coordiF, S_coordiF, diam, diam);
			drawString(j, X_coordiF+10, S_coordiF);
		}
	}

    selectWindow(left_p);
	rename(left_p);
    saveAs("Tiff", saveDir+left_p);
    
    selectWindow(top_p);
    rename(top_p);
    saveAs("Tiff", saveDir+top_p);
    
    selectWindow(n_p);
    rename(n_p);
    saveAs("Tiff", saveDir+n_p);
    
    run("Clear Results");
    run("Close All");	
}


function operation(){
	open(openDir+list[i]);
	n = getTitle();
	run("Reduce Dimensionality...", "slices frames");
	run("Smooth", "stack");
	Stack.getDimensions(width, height, channels, slices, frames);
	for(j=1;j<=frames;j++){
		for(i=1;i<=slices;i++){
			Stack.setPosition(1,i, j);
			run("Find Maxima...", "prominence=50 strict exclude output=[Point Selection]");
			run("Measure");
		}	
	}

// deletion of overlaps
	for (l=getValue("results.count");l>0;l--) {
		Xtent = getResult("X", l-1);
		Ytent = getResult("Y", l-1);
		Xtent_n = getResult("X", l-2);
		Ytent_n = getResult("Y", l-2);
		Slice_c = getResult("Slice",l-1);
		Slice_n = getResult("Slice",l-2);
		Frame_c = getResult("Frame",l-1);
		Frame_n = getResult("Frame",l-2);
		Mean_c = getResult("Mean",l-1);
		Mean_n = getResult("Mean",l-2);
		//if (Slice_c == Slice_n && Frame_c == Frame_n && abs((Xtent^2 + Ytent^2) - (Xtent_n^2 + Ytent_n^2)) < thresh_dist){
		//	Table.deleteRows(l-1, l-1);
		//}
		if (Frame_c == Frame_n && abs((Xtent^2 + Ytent^2) - (Xtent_n^2 + Ytent_n^2)) < thresh_dist){
			if (Mean_c > Mean_n){
			Table.deleteRows(l-2, l-2);}
			if (Mean_c < Mean_n){
			Table.deleteRows(l-1, l-1);}
		}
		
	}

// selection of non-zeros
	for(k=getValue("results.count");k>0;k--){
		//if(getResult("Slice", k-1) == 0){	
		if(getResult("X", k-1) == 15 && getResult("Y", k-1) == 15){
			Table.deleteRows(k-1, k-1);
		}
	}

	newname = n;
	rename(newname);
	saveAs("Results", saveDir+newname+".csv");
	run("Close All");	
}
	
```
