## Supplementary MovieS1-S2, Supplementary Material S1-S2 for "Centrosome-centromere capture range, rather than centrosome arrangement, determines multipolar chromosome segregation pattern after whole-genome duplication": extract_xyz.ijm

```
showMessage("Select Open Folder");
openDir = getDirectory("Choose a Directory");
showMessage("Select Save Folder");
saveDir = getDirectory("Choose a Directory");
list = getFileList(openDir);
thresh_dist = 2;  // parameter for overlapping distance 

for (i=0; i<list.length;i++){
	operation();
};


function operation(){
	open(openDir+list[i]);
	n = getTitle();
	run("Reduce Dimensionality...", "slices frames");
	run("Smooth", "stack");
	Stack.getDimensions(width, height, channels, slices, frames);
	for(j=1;j<=frames;j++){
		for(i=1;i<=slices;i++){
			Stack.setPosition(1,i, j);
			run("Find Maxima...", "prominence=30 strict exclude output=[Point Selection]");
			run("Measure");
			
		}	
	}

// selection of non-zeros
	for(k=getValue("results.count");k>0;k--){
		if(getResult("Slice", k-1) == 0){	
			Table.deleteRows(k-1, k-1);
		}
	}

// deletion of overlaps
	for (l=getValue("results.count");l>0;l--) {
		Xtent = getResult("X", l-1);
		Ytent = getResult("Y", l-1);
		Xtent_n = getResult("X", l-2);
		Ytent_n = getResult("Y", l-2);
		Slice_c = getResult("Slice",l-1);
		Slice_n = getResult("Slice",l-2);
		Frame_c = getResult("Frame",l-1);
		Frame_n = getResult("Frame",l-2);
		if (Slice_c == Slice_n && Frame_c == Frame_n && abs((Xtent^2 + Ytent^2) - (Xtent_n^2 + Ytent_n^2)) < thresh_dist){
			Table.deleteRows(l-1, l-1);
		}
		
	}

	//saveAs("Results", "saveDir");
	newname = n;
	rename(newname);
	saveAs("Results", saveDir+newname+".csv");
	run("Close All");
}

	
```
