## Supplementary MovieS1-S2, Supplementary Material S1-S2 for "Centrosome-centromere capture range, rather than centrosome arrangement, determines multipolar chromosome segregation pattern after whole-genome duplication": README.md

README


Method for obtainig centrosomal coordinate.

1. Select the cell window.
2. Run the ImageJ macro "exact\_xyz.ijm". ・・・(A)
3. Run the ImageJ macro "check\_coordinates.ijm".・・・(B)
4. Add a \[check] column to (A) and record a value of 1 for the corresponding coordinates using (B) as a reference.
5. Run the "220607\_auto\_select\_check1.py" to select correct coordinates
6. Manually sort the coordinates into the four centrosomes based on their relative positions in the preceding and following frames.
