## Supplementary MovieS1-S2, Supplementary Material S1-S2 for "Centrosome-centromere capture range, rather than centrosome arrangement, determines multipolar chromosome segregation pattern after whole-genome duplication": multi_estimate_plane_5_savemass.m

```
%% This program calculate least square plane from xyz dots

%%  parameters for the size of the plane
sizeP_X = 30;
sizeP_Y = 30;

%% for multiple files
FileList = dir('*.csv');
FileNum = size(FileList,1);

%savefolder = "PCA_results";
%mkdir(savefolder);

for i = 1:FileNum 
  
%%  read xyz coordinates
xyz = readmatrix(FileList(i).name);
%save naming
rootname = [FileList(i).name];
csvextension = '.csv';
figextension = '.fig';
%load XYZ;
%DATA = [xyz.X xyz.Y xyz.Slice xyz.Frame];
 %DATA = [xyz.Var1 xyz.Var2 xyz.Var3 xyz.Var4];
 %disp(DATA)
k = 1;
FIRST_FRAME = xyz(1,4);
LAST_FRAME = xyz(end,4);
for j = FIRST_FRAME:LAST_FRAME
    IND(j).A = find(xyz(:,4) == j);
    [NUM ~] = size(IND(j).A);
    IND(j).COORDINATES = xyz(k:k+NUM-1,:);
    COORD = IND(j).COORDINATES;
    k = k + NUM;
    

%%  calculate the center of mass
%Xmass = mean(XYZ(:,1));
%Ymass = mean(XYZ(:,2));
%Zmass = mean(XYZ(:,3));
Xmass = mean(IND(j).COORDINATES(:,1));
Ymass = mean(IND(j).COORDINATES(:,2));
Zmass = mean(IND(j).COORDINATES(:,3));

%%  calculate principle components
XYZ = [IND(j).COORDINATES(:,1), IND(j).COORDINATES(:,2), IND(j).COORDINATES(:,3)];
PCA_XYZ = pca(XYZ);

%% select the third PCA verctor as normal vector
PCA_3X = PCA_XYZ(1,3);
PCA_3Y = PCA_XYZ(2,3);
PCA_3Z = PCA_XYZ(3,3);

%% calculate centrosome-plane distance
Dis1 = abs(PCA_3X*COORD(1,1) + PCA_3Y*COORD(1,2) + PCA_3Z*COORD(1,3) - (PCA_3X*Xmass + PCA_3Y*Ymass + PCA_3Z*Zmass))/sqrt(PCA_3X^2+PCA_3Y^2+PCA_3Z^2);
Dis2 = abs(PCA_3X*COORD(2,1) + PCA_3Y*COORD(2,2) + PCA_3Z*COORD(2,3) - (PCA_3X*Xmass + PCA_3Y*Ymass + PCA_3Z*Zmass))/sqrt(PCA_3X^2+PCA_3Y^2+PCA_3Z^2);
Dis3 = abs(PCA_3X*COORD(3,1) + PCA_3Y*COORD(3,2) + PCA_3Z*COORD(3,3) - (PCA_3X*Xmass + PCA_3Y*Ymass + PCA_3Z*Zmass))/sqrt(PCA_3X^2+PCA_3Y^2+PCA_3Z^2);
Dis4 = abs(PCA_3X*COORD(4,1) + PCA_3Y*COORD(4,2) + PCA_3Z*COORD(4,3) - (PCA_3X*Xmass + PCA_3Y*Ymass + PCA_3Z*Zmass))/sqrt(PCA_3X^2+PCA_3Y^2+PCA_3Z^2);
Dis_Sum = Dis1 + Dis2 + Dis3 + Dis4;


PCA_XYZ = [PCA_3X PCA_3Y PCA_3Z Xmass Ymass Zmass Dis1 Dis2 Dis3 Dis4 Dis_Sum];
%PCA_XYZ = [PCA_3X PCA_3Y PCA_3Z];

csvfilename = [rootname, csvextension];
writematrix(PCA_XYZ, csvfilename, 'WriteMode', 'append')
end

%%  derive the plane
for simX = 1:sizeP_X
  for simY = 1:sizeP_Y
    simZ(simY,simX) = Zmass - (PCA_3X*(simX - Xmass))./PCA_3Z - (PCA_3Y*(simY - Ymass))./PCA_3Z;
  end

  
end

%%  visualize the derived plane and the dots
figure;surf(simZ);hold on;
fig = plot3(IND(LAST_FRAME).COORDINATES(:,1),IND(LAST_FRAME).COORDINATES(:,2),IND(LAST_FRAME).COORDINATES(:,3),'o');
figfilename = [rootname, figextension];
savefig(figfilename) 

%savefig(fig, );
%plot3(XYZ(:,1),XYZ(:,2),XYZ(:,3),'o');
%plot3(Xmass,Ymass,Zmass,'ro');

FILE_OUT = sprintf('NormalVect%d',i);
%FILE_OUT_C = char(FILE_OUT);
save(FILE_OUT,'PCA_XYZ');
close


end
```
